# Membrane constraints reshape synaptotagmin recognition by botulinum neurotoxin B1

**DOI:** 10.1101/2025.08.27.672333

**Authors:** Fodil Azzaz, Oussama El Far, Jacques Fantini

**Author notes:** Corresponding author: Fodil Azzaz, INSERM UA16.

## Abstract

Synaptotagmins 1 and 2 (SYT1 and SYT2) are essential Ca^2^+ sensors in neurotransmission and the functional receptors of botulinum neurotoxin B1 (BoNT/B1). While crystallographic models have defined key contacts, they neglect membrane constraints. Using molecular dynamics simulations in lipid rafts, we uncover how the membrane environment reshaped synaptotagmin conformation and enables critical contacts with BoNT/B1’s lipid-binding loop (LBL). Notably, ganglioside GT1b bridges BoNT/B1 and SYT1, stabilizing their interface through lipid-mediated interactions. This mechanism escapes AlphaFold prediction, which generates non-physiological complexes with steric clashed, revealing a fundamental limitation of current AI methods for membrane-constrained interfaces. Our study demonstrates that lipid rafts create functional binding modes through synergistic protein-lipid interactions, highlighting the epigenetic dimension of protein structure where environment dictates conformation.

## Introduction

Synaptotagmins (SYTs) are synaptic vesicle proteins that function as calcium sensors for neurotransmitter release [1-4]. Synaptotagmin 1 and 2 (SYT1/2) serve as specific protein receptors for botulinum neurotoxin B (BoNT/B), a category A agent that causes botulism by cleaving SNARE (Soluble N-ethylmaleimide-sensitive-factor Attachment protein Receptor) proteins and inhibiting neurotransmission [5-7]. BoNT/B is one of the eight serotypes classified from A to X, of which the serotypes A, B, E and F are associated with human botulism [8-11]. Each serotype is divided into subtypes with different amino acid sequences [12]. Additionally, there are “mosaic BoNTs formed by the combination of the amino acid sequences of different serotypes [13].

The 150 kDa BoNT/B toxin is composed of a light chain (LC, metalloprotease) and a heavy chain (HC) [14, 15]. The HC C-terminal domain (Hcc) mediates receptor binding. X-ray structures of BoNT/B bound to SYT1 or SYT2 fragments have defined the interaction interface within residues 39-49 of SYT1 and 47-58 of SYT2. [16, 17]. In addition to its protein receptor, BoNT/B binds to human gangliosides GT1b and GD1a through a common ganglioside-binding site which is conserved in other toxin types [18-21]. Commonly, gangliosides are packed with cholesterol and sphingolipids in lipid rafts, which are electronegatively charged platforms that attract cationic molecules such as virus proteins or amyloid proteins on the membrane surface [22-28]. Cesare Montecucco proposed a model in which the neurotoxin interacts simultaneously with its protein and glycolipid receptor [29].

However, these structural models present a static and incomplete picture, as they fundamentally lack the native membrane environment. This omission is critically relevant for understanding the function of a unique structural feature of BoNT/B: its lipid-binding loop (LBL, residues E1245 to E1252) [30, 31]. While the LBL is hypothesized to be important for receptor binding, its precise mechanistic role has remained enigmatic. This gap in understanding is highlighted by a key biochemical paradox: although the LBL is dispensable for SYT binding in soluble assays, it is absolutely essential for toxin binding in a membrane context. The current crystallographic data, which show no role for the LBL, are thus insufficient to explain its mandatory function in a physiological setting.

This discrepancy points to a broader limitation: existing models cannot capture the geometric constraints imposed by the lipid bilayer on the conformation of synaptotagmin or the potential synergy between its protein and glycolipid receptors. Therefore, to resolve this paradox and unravel the authentic molecular mechanism of BoNT/B intoxication, there is an urgent need for a structural model that incorporates the toxin in complex with its full repertoire of membrane-embedded receptors within a realistic lipid environment.

The recent revolution in AI-based protein structure prediction offers new opportunities to explore this paradox. We therefore asked whether integrating molecular dynamics with AlphaFold benchmarking could provide the missing structural insights into LBL function in membrane environments.

For this purpose, we present an integrated computational approach combining all-atom molecular dynamics simulations of membrane-inserted complexes with systematic AlphaFold benchmarking. This strategy not only reveals how membrane constraints redefine host-pathogen interactions but also establishes the boundary conditions for AI-based prediction in membrane environments.

## Results

### Membrane constraints reshape synaptotagmin conformation and alter BoNT/B1 binding surfaces

The BoNT/B1-SYT1 and BoNT/B1-SYT2 complexes were inserted into a lipid raft membrane to model the native neuronal environment. To assess the stability of the toxin, we calculated the Root Mean Square Fluctuation (RMSF) of BoNT/B1 in both complexes. The low average RMSF values (1.27 Å for SYT1 and 1.37 Å for SYT2) indicate that BoNT/B1 maintains its overall structural integrity, with most residues displaying a minimal flexibility (< 2 Å) **(Figure 1 A)**.

**Figure 1:**
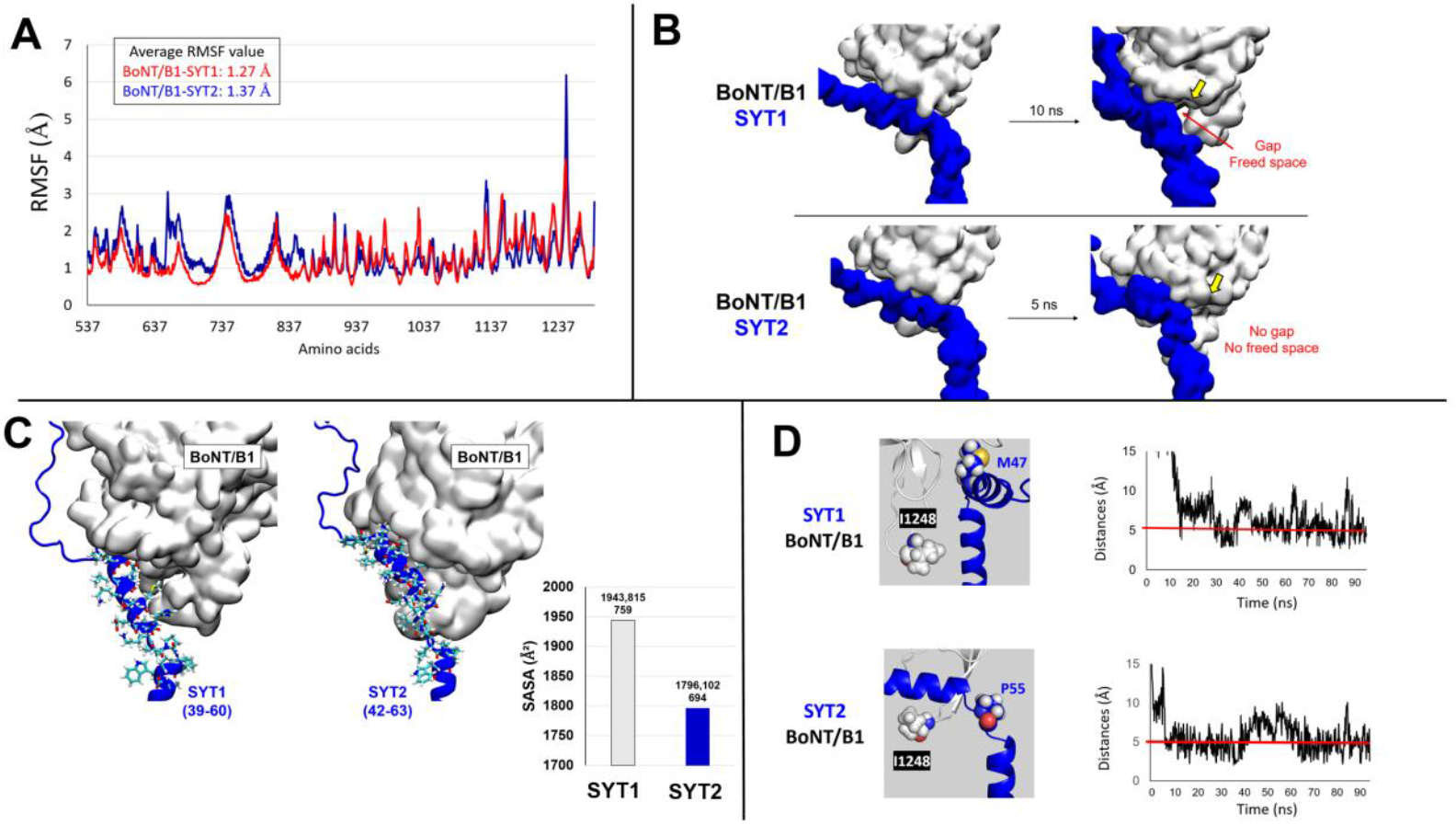
Root Means Square Fluctuation (RMSF) value of BoNT/B1 in complex with SYT1 (red lines) or SYT2 (blue lines). Is framed on top left the graph the average RMSF value for both conditions **(A)**. The membrane environment places stress on the synaptotagmin structure. This reduces the contact surface between SYT1 (upper panel) or SYT2 (lower panel) with BoNT/B1 (highlighted by a yellow arrow). SYT1-2 are depicted as blue surface while BoNT/B1 is shown as a white surface **(B)**. Surface representation of BoNT/B1 (white) in complex with the juxtamembrane domain of SYT1 or SYT2 (residues 42-63). Bar graph showing the average solvent-accessible surface area (SASA) of SYT1 (residues 39-60, white bar) and SYT2 (residues 42-63, blue bar) over time. The observed increase in SASA for the SYT1 complex is consistent with a sustained loss of contact compared to the SYT2 complex. Numerical values represent the mean SASA for each system **(C)**. Snapshots showing residues involved in the interaction between the LBL structure of BoNT/B1 and the SYT1 **(upper panel)** or SYT2 **(lower panel)**. Line graphs on the right side plot the evolution of the minimal distance between the lateral chain of residues I1248-M47 (BoNT-SYT1) and I1248-P55 (BoNT-SYT2) were plotted over time. BoNT/B1 is represented as white cartoon, SYT1 and SYT2 are represented as blue cartoon. The residue I1248 belonging to the LBL structure of BoNT/B1 is depicted as white sphere while the residues M47 of the juxta membrane domain of SYT1 and P55 of the juxta membrane domain of SYT2 are depicted as blue spheres **(D)**

Shortly after the beginning of the simulation, we observed that SYT1 and SYT2 undergo conformational changes due to the constraints imposed by the membrane. As shown in **figure 1 B, upper panel**, SYT1 undergoes stress which, after 10 ns of simulation, forces the conformation of its structure on the side of the ganglioside interaction site of BoNT/B1 (yellow arrow) to move closer to the membrane surface. This movement cancels out intermolecular interactions between the two partners, forming a gap (highlighted by a red arrow). As shown in **figure 1 B, lower panel**, the same phenomenon is observed for SYT2 but interestingly, no gap is formed between both partners. We quantified this loss of contact by calculating the solvent-accessible surface area (SASA) of the synaptotatmin-binding interface on BoNT/B1. The average SASA for the SYT1 complex was significantly larger than that for the SYT2 complex (1944 Å^2^ versus 1796 Å^2^, **figure 1 C**). These preliminary observations suggests a stronger chemical affinity between SYT2 and BoNT/B1 under membrane constraints.

### Membrane constraints enable the lipid-binding loop (LBL) of BoNT/B1 to establish direct contacts with synaptotagmin

Due to the forces that constrained the structure of SYT1 and SYT2 to move closer to the membrane surface, contact points were created between the LBL structure of BoNT/B1 and the synaptotagmins. As shown in **figure 1 D** residues subject to contact are I1248 of the LBL of BoNT/B1 and the residues M47 of SYT1 and P55 of SYT2. Distances between the lateral chain of residues I1248-M47 and I1248-P55 were plotted. The plots reveal that the distances are generally below 5 Å throughout the simulation, indicating that the residues are often in contact together via van der Waals interactions.

### GT1b ganglioside bridges BoNT/B1 and SYT1, stabilizing the membrane-induced complex

As shown in **figure 1 B and figure 1 b**, a gap is formed between SYT1 and BoNT/B1 due to the membrane constraints while such a gap is not formed between SYT2 and BoNT/B. In response to this gap, the simulation predicts that the GT1b molecule that was initially mainly interacting with the aromatic residue W1262 of BoNT/B1 (**figure 2 A**), adapts its conformation to end up interacting with the cationic residues K1187 and K1188 of BoNT/B1 and the aromatic residue H52 of SYT1 (**figure 2 B, C, D, E and F**). The GT1b molecule appears to act as a bridge between BoNT/B1 and SYT1 (**figure 2 F**) that stabilizes the link between the toxin and its protein receptor by occupying the space generated by the membrane constraints. This ganglioside bridging illustrates a synergy between protein and glycolipid receptors that cannot be captured without membrane context.

**Figure 2:**
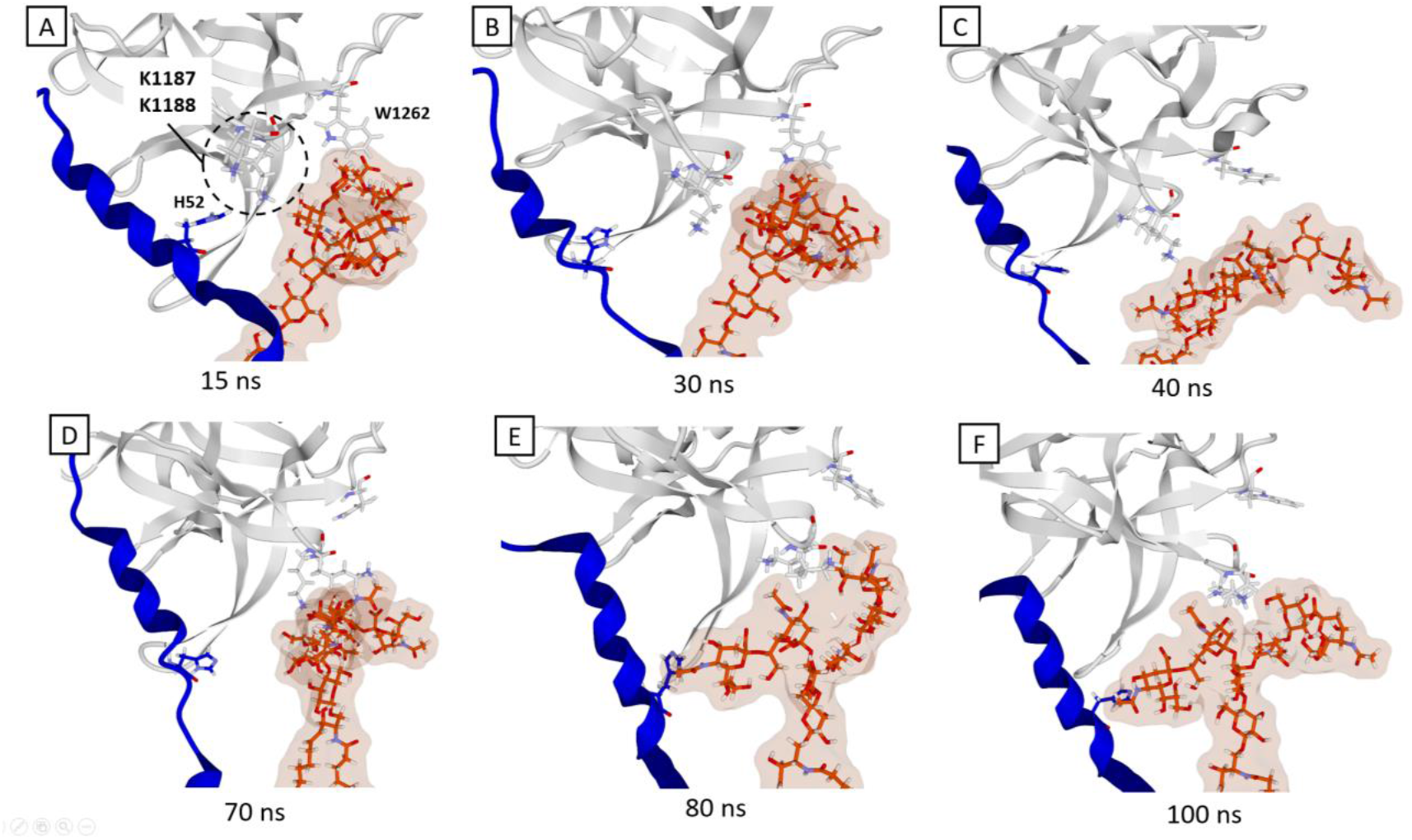
GT1b ganglioside bridges BoNT/B1 and SYT1. Snapshots taken at 15, 30, 40, 70, 80, 100 ns showing the panel of conformations adopted by the gangliosides and its insertion in the SBP (**A, B, C, D, E and F**). The ganglioside is in orange, h-SYT1 in blue and the toxin in white.

### A lipid raft-associated GT1b molecule interacts with the LBL of BoNT/B

In addition to SYT1 and SYT2, the simulation suggests that the LBL structure of BoNT/B1 also interacts with a lipid raft-associated GT1b molecule. The GT1b molecule interacts with this structure mainly via residues R1242 and F1250. Snapshots showing the kinetics of the interaction mechanism between the ganglioside and the lipid-binding loop are shown in **figure 3**. These snapshots show that, initially and after 10 ns of simulation, the sugar moiety of the ganglioside is in a closed conformation able to interact only with the aromatic residue F1250. After 35 ns, the polar part of the ganglioside opens, allowing simultaneous interaction with residues R1242 and F1250, as shown in the snapshot obtained after 68 ns of simulation.

**Figure 3:**
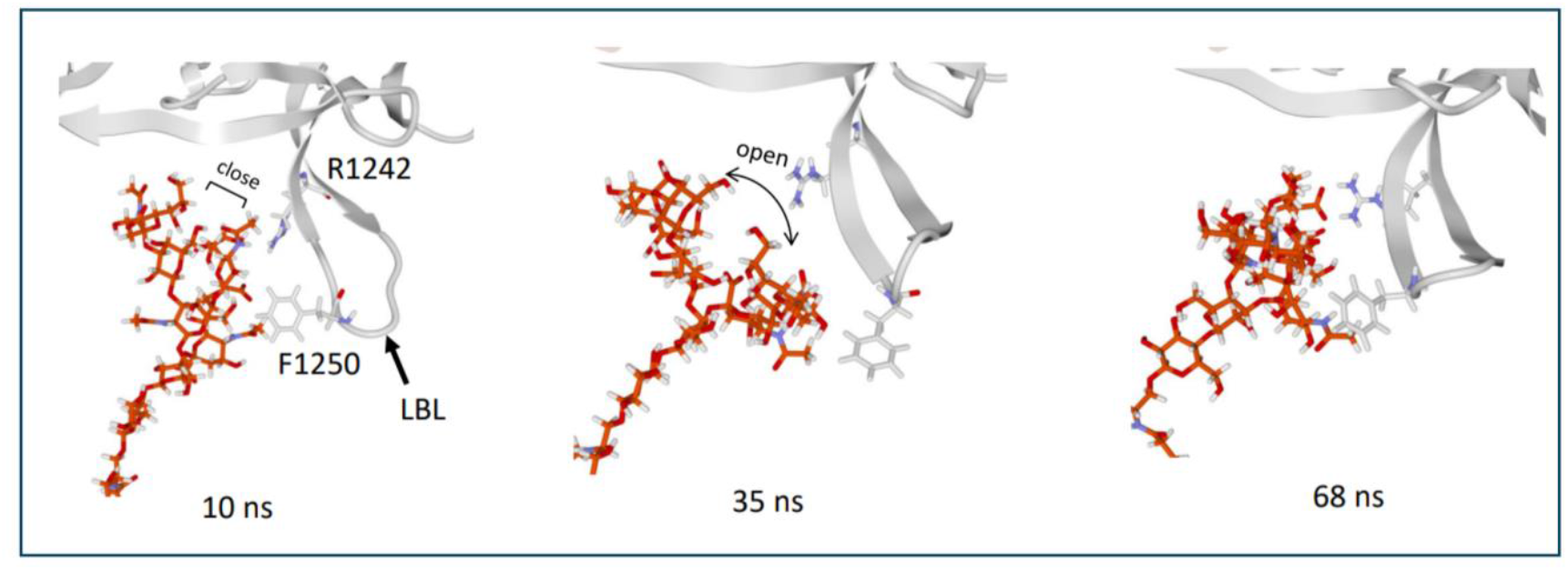
Snapshots of the interaction interface between the ganglioside and the lipid-binding loop of BoNT/B. Kinetics of the interaction mechanism of a GT1b molecule with the lipid-binding loop of BoNT/B1 at times 10, 35 and 68 ns. The ganglioside is shown as orange sticks, BoNT/B1 is shown as a white cartoon and residues R1242 and F1250 of BoNT/B1 are represented as white sticks.

### AlphaFold-Multimer fails to predict a realistic membrane-constrained complexes

To assess whether current AI methods could capture our identified mechanism, we benchmarked AlphaFold-Multimer on BoNT/B-SYT complexes. When using structural templates from PDB, AlphaFold correctly localized synaptotagmin to its binding site but showed inconsistent confidence in functionally critical regions: the juxtamembrane domain displayed mixed high/low confidence for SYT4, while the LBL showed low confidence for SYT1 and medium confidence for SYT2 **(figure 4 A and B)**.

**Figure 4:**
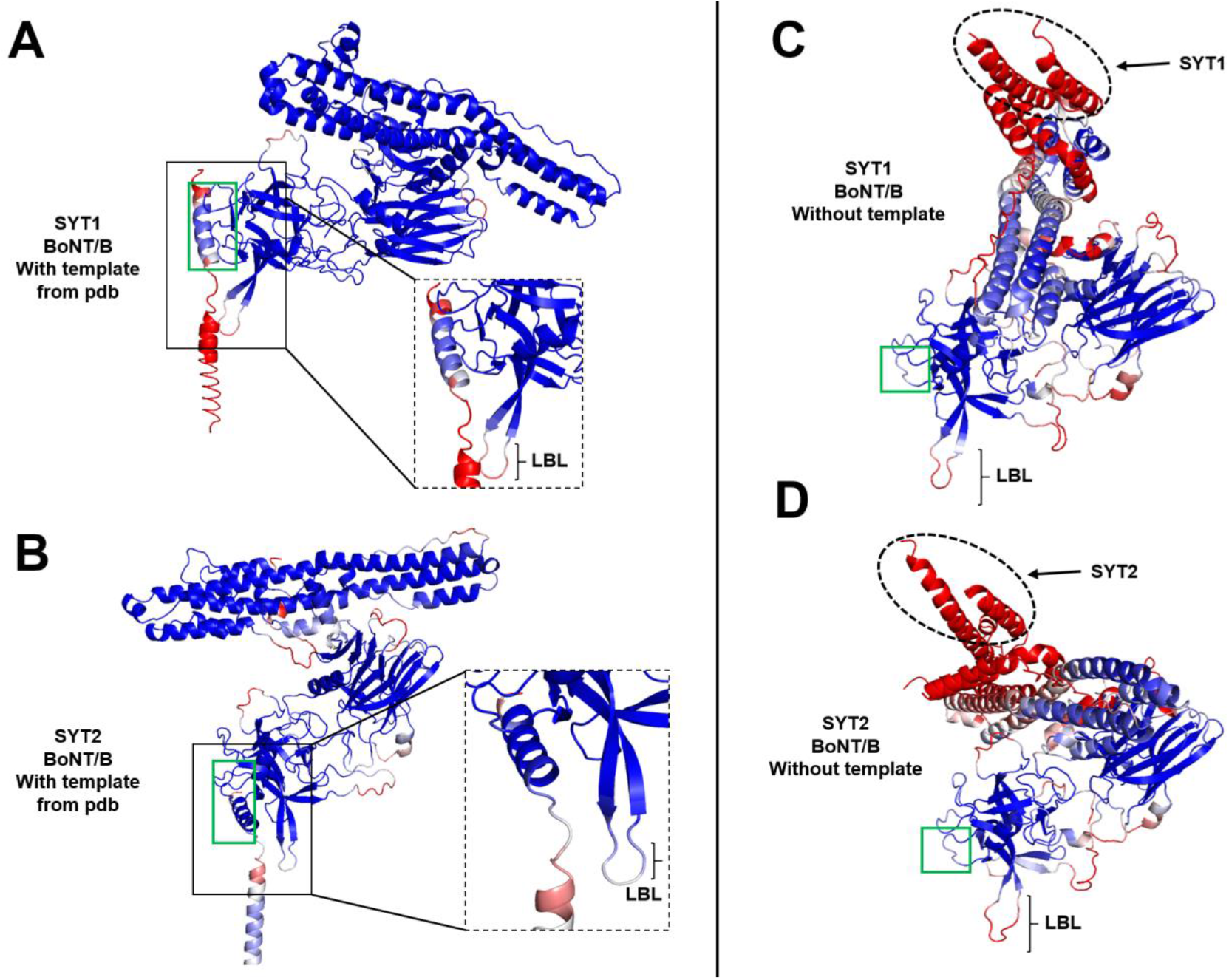
Prediction of BoNT/B-SYT complex using AlphaFold-Multimer. Predicted structure of BoNT/B in complex with SYT1 **(A, C)** or SYT2 **(B, D)** generated with **(A, B)** or without **(C, D)** structural templates from PDB. Structures are colored by per-residue confidence (pLDDT), with blue indicating high confidence, white medium and red low confidence. The lipid-binding loop (LBL) is indicated by braces and the synaptotagmin-binding interface is outlined in green, SYT1 and SYT2 are highlighted by a dashed oval circle **(C, D)**. Note the low confidence in membrane-proximal regions and the LBL across all predictions.

More strikingly, template-free predictions revealed fundamental limitations. AlphaFold placed synaptotagmin in non-physiological positions, with erroneous interactions between synaptotagmin and the translocation domain of BoNT/B instead of the correct binding site **(figure 4 C and D)**. These models also exhibited uniformly low confidence for synaptotagmin regions, indicating the inability of AlphaFold to confidently resolve membrane-proximal interfaces without structural priors for BoNT/B-SYT complexes.

Structural analysis of the predicted BoNT/B-synaptotagmin complexes revealed severe molecular incompatibilities. Surprisingly, complexes involving SYT1 (generated with templates) and both SYT1/2 (generated without templates) exhibited significant steric clashes **(figure 5 A-C)**. Interaction energy calculations confirmed these were not minor artifact but severe structural violations, with clash energies exceeding 200 kJ/mol for complexes involving SYT1 **(figure 5 A and B)** and 50 kJ/mol for complexe involving SYT2 **(figure 5 C)**.

**Figure 5:**
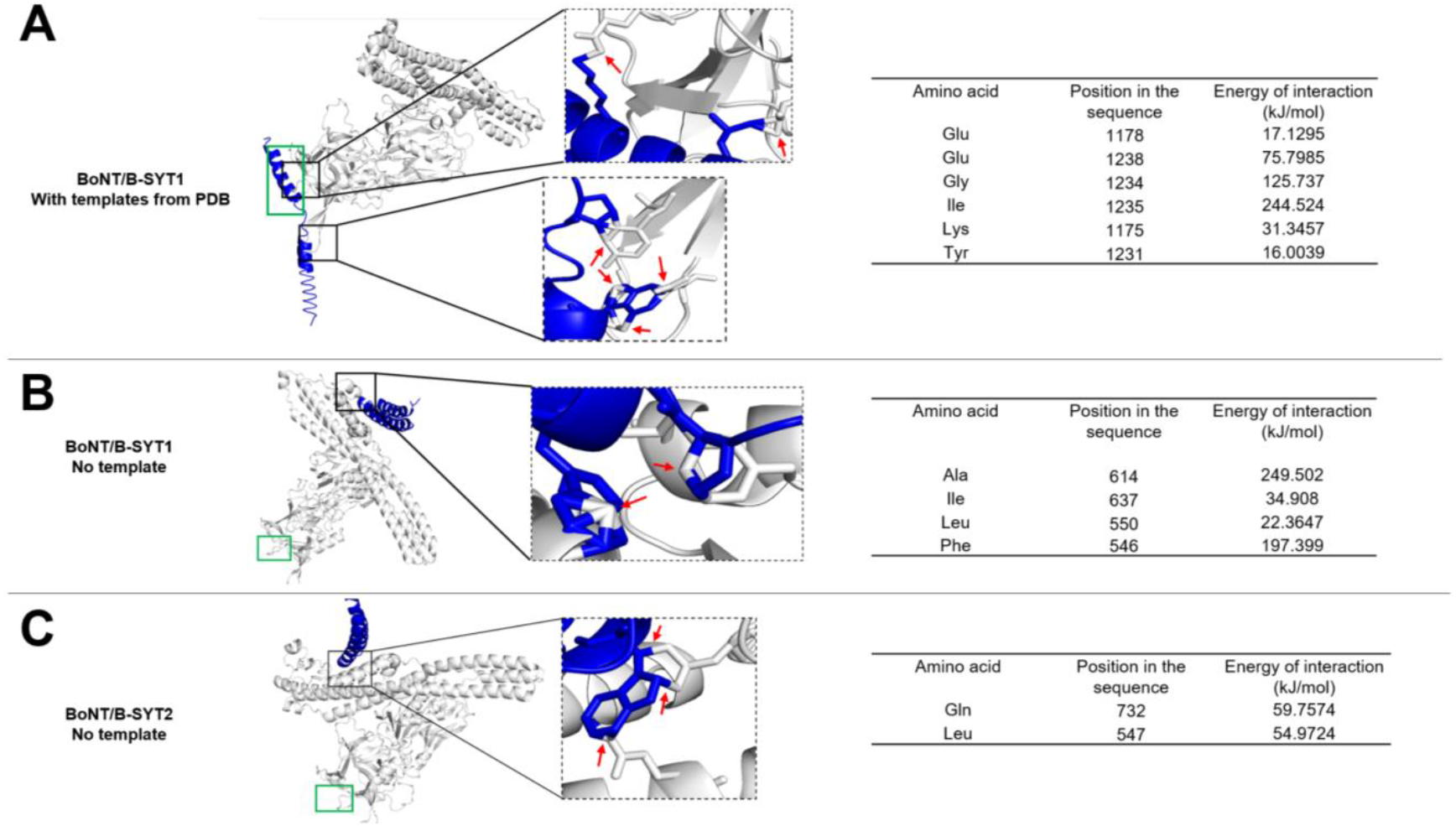
AlphaFold-Multimer generates BoNT/B-synaptotagmin complexes with steric clashes. Predicted structure of BoNT/B in complex with SYT1 **(A, B)** or SYT2 **(C)** generated with **(A)** or without **(B, C)** structural templates from PDB. For each structure is shown a zoom revealing the molecular details of steric clashes highlight with red arrows. Each structure is linked to a table of energy showing the toxin’s residues involved in important steric clashes. Synaptotagmin binding sites is outlined in green.

At the molecular level, we observed toxin residues unrealistically integrated within the aromatic rings of synaptotagmin, a physically impossible arrangement that fundamentally disrupts the protein architecture and precludes any meaningful biological interpretation or further modeling. These steric clashes do not represent minor imperfection, but rather, physically impossible errors that fundamentally invalidate the predicted models. Atoms cannot occupy the same space that aromatic rings, such predictions violate the most basic principles of structural chemistry.

The BoNT/B-SYT2 complex generated with PDB templates was the only AlphaFold prediction lacking steric clashes. We therefore assessed whether this structurally model was nonetheless compatible with a membrane environment. Comparative analysis with our simulated structure revealed a critical discrepancy. The juxtamembrane domain of the predicted SYT2 was positioned abnormally close to the membrane surface **(figure 6 A-E)**. This constitutes a major functional problem. Our previous work established that GT1b specifically interacts with the juxtamembrane fomain of synaptotagmins via a defined ganglioside-binding site [32]. The compressed conformation of AlphaFold’s SYT2 sterically occludes this site, making specific interaction with GT1b impossible **(figure 6 D and E)**. Thus, even AlphaFold’s most plausible prediction fails to capture the membrane-compatible geometry required for physiological function.

**Figure 6:**
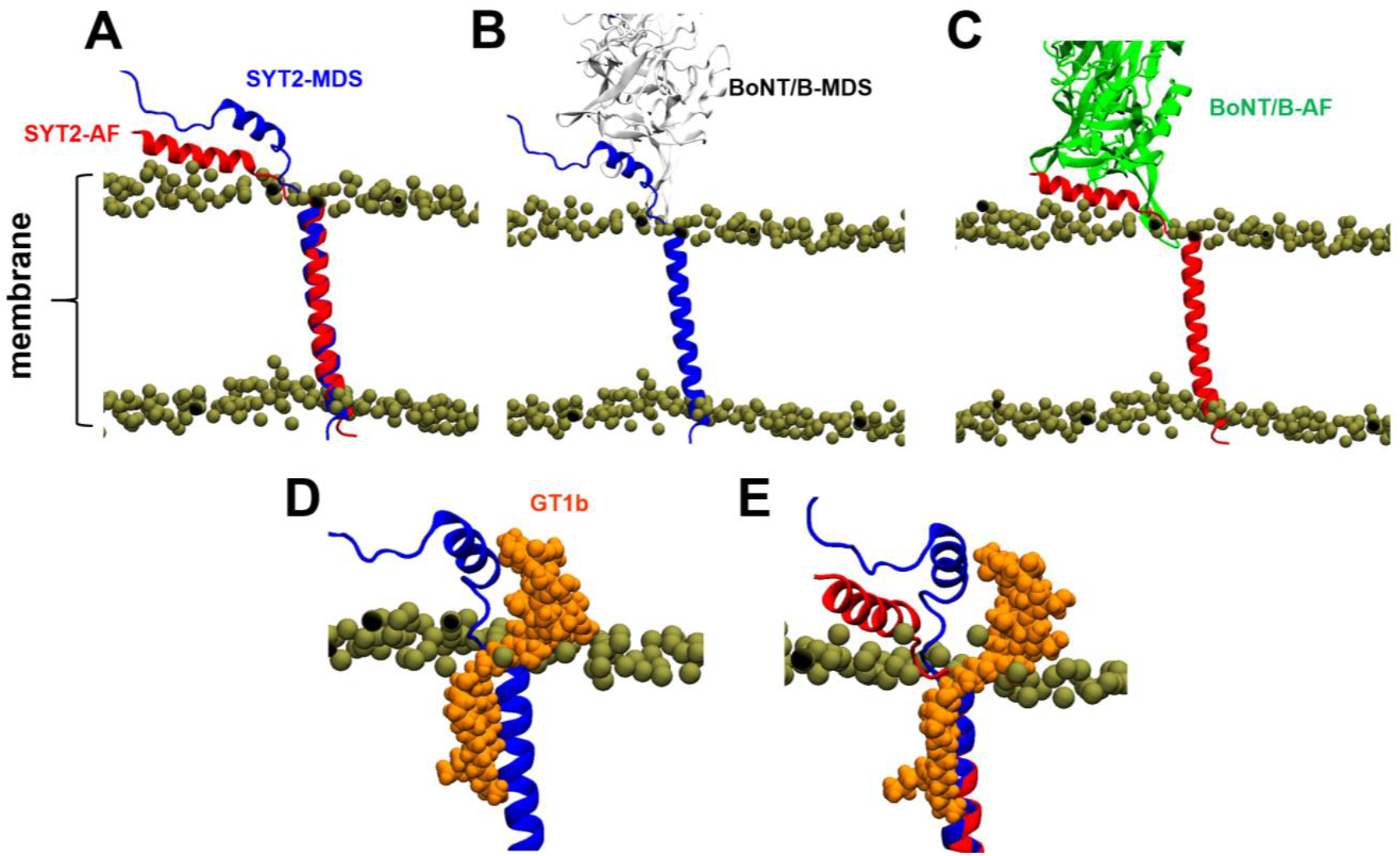
AlphaFold-Multimer complexes are incompatible with membrane constraints. Insertion of the predicted complex in membrane based on the alignment of the transmembrane helix of predicted SYT2 “AF” in red with simulated SYT2 “MDS” in blue **(A)**. Overview of simulated **(B)** or predicted **(C)** BoNT/B-SYT2 complex in the membrane. Snapshot showing a GT1b molecule (orange) interacting with the juxta membrane domain of simulated SYT2 **(D)**. Same as **(D)**, but predicted SYT2 is also displayed. In each image, the phosphorus atom of POPC molecules is depicted as a brown sphere. Phosphorus atom is the frontier between the polar and apolar part of lipids. Note that, the juxta membrane domain of predicted SYT2 is closer to the membrane surface, making it non possible for GT1b to comfortably interact with.

Collectively, these results demonstrate that AlphaFold fails to predict the BoNT/B-synaptotagmin interface across multiple critical dimensions. It produces either structurally impossible complexes with steric clashes, or topologically incorrect models that are incompatible with membrane constraints and essential lipid interactions, revealing a fundamental limitation in modeling membrane-driven allostery.

## Discussion

Our MDS provide the first atomic-resolution model of the BoNT/B1 receptor complex within a realistic lipid raft environment, revealing how membrane constraints and gangliosides collaboratively reshape the toxin’s interaction with its synaptotagmin receptors. While crystallographic studies have been instrumental in identifying binding pockets, their soluble nature inherently overlooks the profound impact of membrane topology on synaptotagmin conformation—a limitation particularly relevant given the established direct interaction between ganglioside and the synaptotagmin juxtamembrane domain via ganglioside-binding site (GBS) motif absent in crystal structures [32, 33].

A key prediction of our study is that the lipid raft environment imposes distinct geometric constraints on SYT1 and SYT2. Notably, SYT1 is more severely impacted, resulting in a discernible gap at its interface with BoNT/B1 (**figure 1 B and C**). We propose that this membrane-induced destabilization of the SYT1 complex is compensated by the dynamic recruitment of a GT1b molecule, which moves from its initial interaction with W1262 to bridge the freed space, engaging both BoNT/B1 (K1187 + K1188) and SYT1 (H52) (**Figure 2**). This provides a compelling mechanistic explanation for the long-standing observation that BoNT/B1 binding to SYT1 is strictly ganglioside-dependent [34].

Furthermore, our simulations reveal a novel and functionally critical interaction: the direct contact between the juxtamembrane domains of both SYT1 and SYT2 with the BoNT/B1 lipid-binding loop (LBL) (**Figure 1 D**). This finding resolves a significant paradox in the field. As demonstrated by Stern et al., the LBL is dispensable for SYT2 binding in soluble assays but becomes essential in a membrane environment [31]. Our model accounts for this discrepancy directly; in the context of membrane constraints, the LBL serves as a crucial docking site for synaptotagmin, a role that is unnecessary and thus invisible in soluble structures (**Figure 7**).

**Figure 7:**
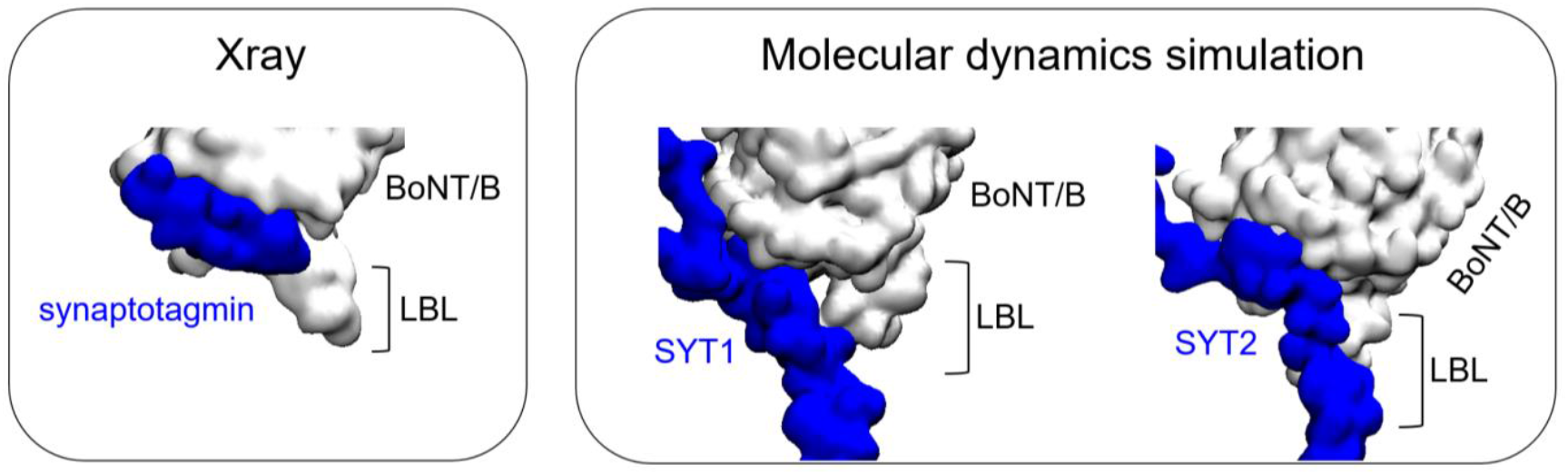
Snapshot showing the model of synaptotagmin in complex with BoNT/B1 in a liquid environment. These crystallographic data are obtained in the PDB (entry 6G5G) **(left panel)**. Snapshots showing the model of SYT1 or SYT2 in complex with BoNT/B1 in a membrane environment **(right panel)**. BoNT/B is represented as white surface while the synaptotagmin is represented as blue surface.

A principal strength of our molecular dynamics approach is its capacity to resolve atomic-level dynamics that are presently beyond the reach of direct experimental observation. While techniques like cryo-electron microscopy provide indispensable static snapshots, they cannot capture the real-time, lipid-dependent conformational rearrangements that are central to the BoNT/B1 recognition mechanism. The dynamic formation of the GT1b bridge, the membrane-induced contact with the LBL, and the differential stability of the SYT1 and SYT2 complexes are paradigm examples of such transient yet functionally critical events. Thus, our simulations provide a vital functional complement to structural databases, offering a mechanistic narrative for biochemical observations that have until now lacked a structural rationale.

The functional importance of the LBL is further underscored by its role in ganglioside recognition. We observed a lipid raft-associated GT1b molecule engaging the LBL through residues R1242 and F1250 (**Figure 3**). This is in full agreement with experimental data showing that LBL deletion or a F1250R point mutation abolishes BoNT/B binding to gangliosides in nanodiscs [31], validating our model’s predictive power.

Strikingly, our benchmarking with AlphaFold-Multimer reveals that this membrane-constrained mechanism completely escapes AI-based prediction. AlphaFold not only fails to recapitulate the lipid-bridging interface and the critical LBL-synaptotagmin contacts we identifies but also generates structurally unrealistic complexes with steric clashes and topologically unrealistic membrane orientations. This fundamental limitation demonstrates that current AI methods cannot capture environmentally-induced conformational changes.

In conclusion, our work moves beyond static models to demonstrate that the membrane is an active participant in forming the BoNT/B1 receptor complex. From a fundamental point of view, our study reinforces the concept according to which the structure of an initially disordered domain of the juxtamembrane part of synaptotagmin can be controlled by different ligands that can act in synergy, particularly when these ligands are part of a lipid raft. This particularity of intrinsically disordered proteins (IDPs) is an unexpected refinement of biology which is totally ignored by an artificial intelligence such as AlphaFold which only considers the amino acid sequence to predict 3D structures of proteins. This algorithm lacks the consideration of IDP ligands and more broadly all the environmental parameters that we have grouped together in the concept of Epigenetic Dimension of Protein Structure (EDPS). This new theory of protein structure has already made it possible to highlight the role of raft lipids (gangliosides and cholesterol) in the propagation of transmembrane conformational waves initiated by the binding of a ligand to its receptor. Taking into account this conformational flexibility inherent to the intrinsic disorder of critical receptor domains will make it possible in the future to develop innovative therapeutic approaches that are less “frozen in stone” than those suggested by classic models [35, 36]. Furthermore, our findings suggest that the role of membrane-induced geometric constraints and lipid bridging is a general mechanism that must be considered for any protein-protein interaction occurring at the membrane surface. This study serves as a critical warning against extrapolating the mechanisms of membrane protein interactions solely from structures solved in soluble, non-membrane environments. It demonstrates that the true functional binding mode can be entirely reshaped by the lipid bilayer.

## Methods

### System set-up

The initial coordinates of ganglioside and cholesterol molecules composing the lipid raft were obtained using the glycolipid modeler tool and bilayer builder available on charmm-gui [37]. The structure of BoNT/B1 (from 537 to 1291) was obtained from PDB : 2NP0 and the missing regions were built with SWISS-MODEL [38]. The luminal domain of SYT1 and SYT2 were made de-novo using HyperChem [39]. A molecular docking of the complex BoNT/B-SYT1 or 2 were obtained with HyperChem in a way that respects the geometric constraints of membrane topology. The glycosylations of SYT1and SYT2 were added using the tool PDB reader of charmm-GUI [40] [41]. An energy minimization of the complexes was performed with the Polak-Ribière conjugate gradient algorithm, with the Bio+(Charmm) force field in Hyperchem and a root-mean-square (RMS) gradient of 0.01 kcal. Å^−1^.mol^-1^. Finally, the complexes and the lipid raft were placed in a lipid bilayer that mimics the lipid composition of neural membranes (outer leaflet: 50% POPC and 50% cholesterol | inner leaflet: 30% POPS/20% POPE and 50% cholesterol) [42]. Then, the systems were solvated with TIP3P water and neutralized with the addition of Na^+^ and Cl^-^ions at a final concentration of 0.15 mol/L using the tools “Solvation” and “add ions” available on VMD [43].

### Molecular dynamics simulations

The system was simulated using NAMD 2.14 for Windowsx64 [44] coupled with the all-atom Force Field Charmm36m which has been improved for the calculation of IDPs [45]. The cutoff for the calculation of non-bonded interaction was set to 12 Å. The PME calculation algorithm was used to calculate long range electrostatic interactions. The covalent bonds involving hydrogen atoms were constrained using the algorithm SHAKE. First, the whole systems were minimized for 20000 steps. The apolar parts of the lipids were melted for 1 ns at 310K in the NVT ensemble to get rid of the crystalline organization of the lipid bilayer. Then, the systems were equilibrated at constant temperature (310 K) and constant pressure (1 atm) during 15 ns with constraint on protein followed by 15 ns of equilibration with constraint on backbone only. Finally, the non-constrained runs were performed at constant temperature (310 K) and constant pressure (1 atm) for 100 ns with a time step of 2 fs. The coordinates of the system evolution were backed up every 0.1 ns corresponding to 50000 steps

### AlphaFold-Multimer Predictions

BoNT/B-synaptotagmin complex structures were predicted using AlphaFold-Multimer implemented through ColabFold v1.5.5 (https://colab.research.google.com/github/sokrypton/ColabFold/blob/main/AlphaFold2.ipynb). Sequences of BoNT/B1 (residues 537-1291) and SYT1 (residues 32-87) or SYT2 (residues 40-95) were submitted. Predictions were performed with and without structural templates using default parameters. For each complex, only the rank 1 complex as been studied.

### Structural analysis and validation

Predicted models were analyzed using PyMoL. Per-residue confidence scores (pLDDT) were visualized using the spectrum command with the ‘blue_white_red’ palette (range: 50-90). Interaction energies for each predicted complexes were calculated using Molegro Molecular Viewer to quantify structural stability.

